# LNS8801: An enantiomerically pure agonist of the G protein-coupled estrogen receptor suitable for clinical development

**DOI:** 10.1101/2024.11.26.625421

**Authors:** Christopher A. Natale, Sophia Mercado, Richard Zhuang, Cristina Aguirre-Portolés, Israel Olayide, Christopher K. Arnatt, John T. Seykora, Tina K. Garyantes, Wayne Luke, Todd W. Ridky

## Abstract

Estrogen effects in tissue are mediated in part through activation of the surface estrogen receptor GPER, a broadly expressed G protein-coupled receptor that impacts a wide range of normal and pathologic processes, including metabolism, vascular health, inflammation, and cancer. A commonly used synthetic and specific GPER agonist, named G-1, antagonizes tumors by promoting cellular differentiation and enhancing tumor immunogenicity. G-1 is a racemic compound, and since its discovery, the question of whether both enantiomers display agonist activity or the agonist activity resides primarily in a single enantiomer has never been fully resolved. Herein, we disclose the isolation of the pure enantiomers of G-1 and determine that the desirable activity resides exclusively in 1 enantiomer, named LNS8801, whose configuration we have unambiguously determined by single crystal x-ray structure analysis. Using preclinical models, we show that LNS8801 suppresses cancer in a GPER-dependent manner and that LNS8801 is efficacious when administered orally. Further, we show that GPER is widely, but not ubiquitously, expressed in both normal and malignant human tissues. In addition, an attenuated response to LNS8801 is observed in a common germline coding variant in human GPER. These findings support ongoing human cancer trials with LNS8801 and suggest that the germline GPER genotype may serve as a predictive biomarker of therapeutic response.

## Introduction

GPER (G protein-coupled estrogen receptor) is a transmembrane G protein-coupled receptor (GPCR) activated by endogenous estrogens. Unlike the classical nuclear estrogen receptors, GPER does not use a genomic mechanism for its actions but rather directly activates rapid G protein signaling. The relative lack of GPER signaling, particularly in biological males and postmenopausal women, has been implicated in various diseases, including cancer, metabolic disorders, and cardiovascular disease, making GPER a target for agonist drug development (1,2). Recent research has determined that GPER signaling is tumor-suppressive in multiple tissue settings (3,4). Although no approved drugs target GPER, GPCRs are generally “druggable” as over 40% of all drugs approved by the U.S. Food and Drug Administration modulate GPCRs (5).

The rationale for targeting GPER in cancer stems from the epidemiological observation that female sex and history of pregnancy are associated with decreased incidence of and improved stage-specific survival for many common cancers, including melanoma, pancreatic ductal adenocarcinoma (PDAC), non–small cell lung cancer, leukemias, and colon carcinomas, among others (6–13). Although the mechanism underlying this protective effect has remained unknown since it was first appreciated over 50 years ago (14), we considered the possibility that signaling induced by female sex hormones may protect against some malignancies. We hypothesized that understanding the mechanisms responsible for the apparent female protective effect could lead to the identification of new therapeutic targets for cancer, including malignancies that are not classically considered sex hormone responsive.

We recently demonstrated that nonclassical estrogen signaling through the GPER inhibits melanoma and PDAC through similar mechanisms, driving cellular differentiation and inhibition of proliferation (3,4). In mice with therapy-resistant syngeneic tumors, we discovered that systemic administration of a specific small molecule GPER agonist (15) induces differentiation in tumor cells, inhibits their proliferation, and simultaneously renders the tumors more immunogenic. Combination therapy with a GPER agonist and an anti–programmed death-1 (PD-1) immune checkpoint inhibitor demonstrated synergistic antitumor activity, with a high percentage of complete responses that were associated with long-lasting tumor immunity. This immunity protected the mice against subsequent rechallenge with treatment-naive tumor cells. GPER is expressed in many tissues, (16) and signaling downstream of GPER is mediated by ubiquitous cellular proteins. The use of common cellular machinery for signaling likely explains why many cancers are similarly inhibited by GPER agonists in preclinical models.

Consistent with our data in melanoma and PDAC, others have demonstrated that GPER signaling is tumor-suppressive in cancers that are not traditionally considered hormone responsive, including lung cancer, colon cancer, and melanoma (17–19). These preclinical data, coupled with data here showing GPER expression in a wide range of normal and malignant human tissues, support the idea that GPER may be a therapeutically useful target for a wide range of cancer types.

Since its first discovery as a GPER against, the racemic compound G-1 has been used in virtually all subsequent studies exploring selective activation of GPER as it is readily available from a variety of chemical suppliers. G-1 is structurally dissimilar to estradiol, the endogenous GPER agonist. Racemic G-1 is selective for GPER as binding studies show no binding to nuclear estrogen receptors (15) or 25 other medically relevant GPCRs, even at high concentrations (>10 **μ**M) (39). In the literature, G-1 (15) is often assumed to be an enantiomerical pure molecule rather than a racemic compound, even though the reported synthesis of G-1 affords a racemic product and the name G-1 is assigned to a racemic compound. In addition, the GPER agonist G-1 that is sold by a variety of chemical supply houses is a racemic compound. The enantiomers of racemic G-1 do not interconvert under physiological conditions. While the original work identifying racemic G-1 as a GPER agonist recognized that the enantiomers of G-1 could exhibit differing activities, this question was never fully and unambiguously addressed. Accurately determining the activity of each of the enantiomers of racemic G-1 is critically important to interpret the results of preclinical studies and translate this work into a clinically useful human therapeutic. Using preparative chiral chromatography, we have successfully isolated and purified multigram quantities of each of the enantiomers in racemic G-1. In addition, we have determined the absolute configuration of the active enantiomer called LNS8801 (1-[(3aS,4R,9bR)-4-(6-bromo-1,3-benzodioxol-5-yl)-3a,4,5,9b-tetrahydro-3H-cyclopenta[c]quinolin-8-yl] ethanone) by single crystal x-ray structural determination (21).

We report on experiments that show that only 1 enantiomer exhibits the desirable biological activity exhibited in previous reports in the literature that used racemic G-1. We demonstrate that LNS8801 is orally efficacious at low concentrations in a murine model, that the anticancer activity of LNS8801 is dependent on GPER, and that the effect of LNS8801 is attenuated by a naturally occurring common germline GPER variation in humans (rs11544331 Pro16Leu). Additonally, we characterized the GPER protein expression in a variety of normal tissues and select malignacies.

## Results

### LNS8801 is the active enantiomer in racemic G-1

The synthesis of G-1 (15,22) results in a racemic mixture primarily composed of 2 enantiomers (**Figure 1A**), LNS8801 and LNS8812. The 2 enantiomers present in G-1 were isolated using chiral chromatography, resulting in LNS8801 and LNS8812 (21). The activity of LNS8801 and LNS8812 were evaluated in vitro using multiple models that had previously been demonstrated to be susceptible to GPER-mediated suppression, including YUMM1.7 murine melanoma and 2838c3 murine PDAC (3,4). G-1 inhibited the proliferation of both cell lines as expected. LNS8801 was more potent than G-1, while LNS8812 and vehicle had no measurable effect (**Figure 1B-C**). Effects of LNS8801 were validated by measuring cAMP, which is an established readout for GPER and Gs signaling (23,24). We treated human promyelocytic leukemia (HL-60) cells with vehicle, G-1, LNS8801, and LNS8812 and observed cAMP inductions with G-1 and LNS8801 treatment but not vehicle or LNS8812 treatment (**Supplemental Figure 1A**).

**Figure 1:**
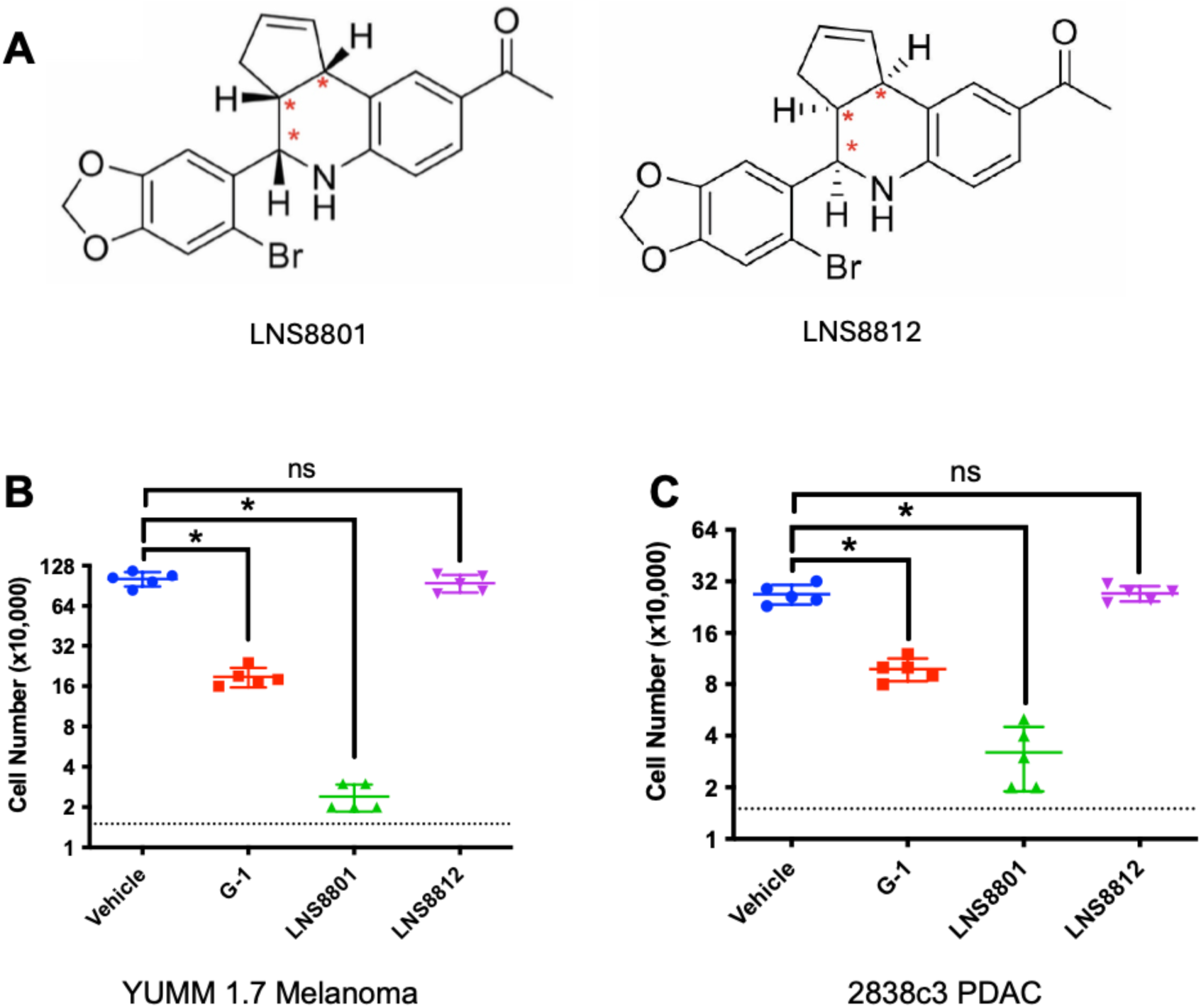
G-1 is a racemic compound. (A) Chemical structures of the enantiomers of racemic G-1, LNS8801 and LNS8812. (B) Proliferation of YUMM1.7 melanoma cells treated with 500 nM G-1, LNS8801, or LNS8812. n=5 per group, * denotes significance by one-way ANOVA, alpha = 0.05. (D) Proliferation of 2838c3 PDAC cells treated with 500 nM G-1, LNS8801, or LNS8812. n = 5 per group, * denotes significance by one-way ANOVA, alpha = 0.05. The dotted line indicates the number of cells in each well at the start of the 4-day incubation.

The activities of each molecule were confirmed in vivo by subcutaneously injecting G-1, LNS8801, and LNS8812 in vivo in the murine 2838c3 PDAC tumor model. G-1 had moderate antitumor effects as expected, while LNS8801 had significantly more activity, including tumor clearance (**Figure 2A-B**). LNS8812 and vehicle lacked any measurable effect on tumor growth. LNS8801 had significantly improved activity compared with G-1 (**Figure 2B)**.

**Figure 2:**
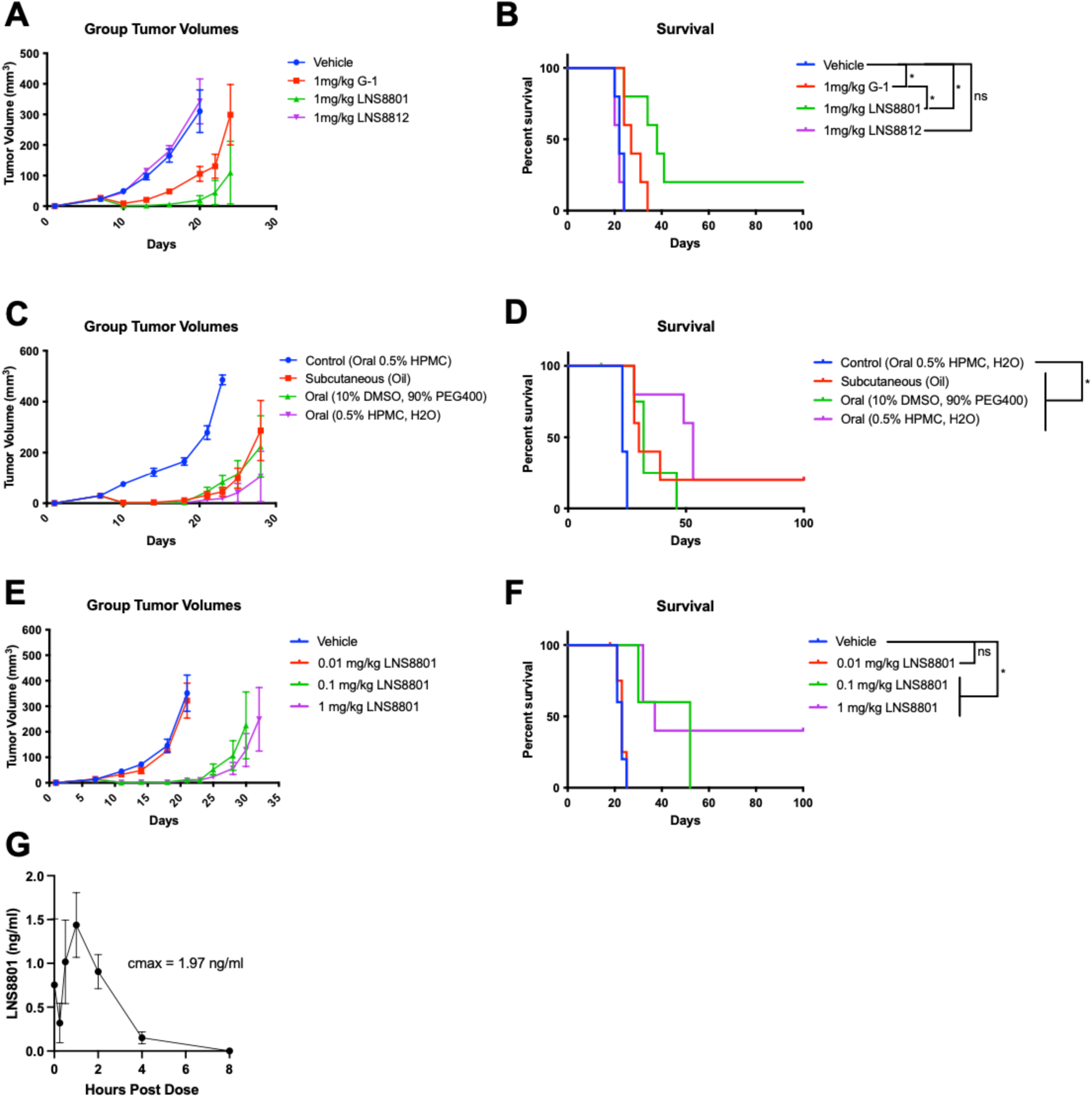
LNS8801 is orally active in vivo. (A) Tumor volumes and (B) Kaplan-Meier survival curves measured over time in 2838c3 PDAC-bearing mice treated with subcutaneously delivered vehicle (sesame oil), G-1, LNS8801, and LNS8812 at 1 mg/kg, significance by log-rank (Mantel-Cox). (C) Tumor volumes and (D) Kaplan-Meier survival curves measured over time in 2838c3 PDAC-bearing mice treated with control vehicle (0.5% HPMC, oral) or LNS8801 delivered subcutaneously in oil, orally solubilized in 10% DMSO/90% PEG400, or orally suspended in 0.5% HPMC at 1 mg/kg doses, significance by log-rank (Mantel-Cox). (E) Tumor volumes and (F) Kaplan-Meier survival curves measured over time of 2838c3 PDAC-bearing mice treated orally with vehicle, 0.01, 0.1 or 1 mg/kg LNS8801, significance by log-rank (Mantel-Cox). (G) Pharmacokinetic analysis of mice treated with oral 0.1 mg/kg LNS8801.

We determined that LNS8801 was efficacious when delivered either subcutaneously or orally (**Figure 2C-D**). In a dose-ranging study, the antitumor effects of LNS8801 saturated at 0.1 mg/kg (**Figure 2E-F**). We performed a pharmacokinetic analysis of mice treated for 3 consecutive days at 0.1 mg/kg and observed a maximum plasma concentration of 1.29 ng/mL on the third day, which corresponds to a 4.8-nM peak plasma exposure (**Figure 2G).** Antitumor efficacy at this exposure is consistent with the low nanomolar binding affinities of endogenous estrogens and selective agonists on GPER (15).

Using modeling (molecular mechanics energies combined with generalized Born and surface area continuum solvation, MM-GBSA), we calculated the binding energies of both enantiomers to GPER and determined that the LNS8801-GPER complex has lower free energy than the LNS8812 complex (**Figure 3**).

**Figure 3:**
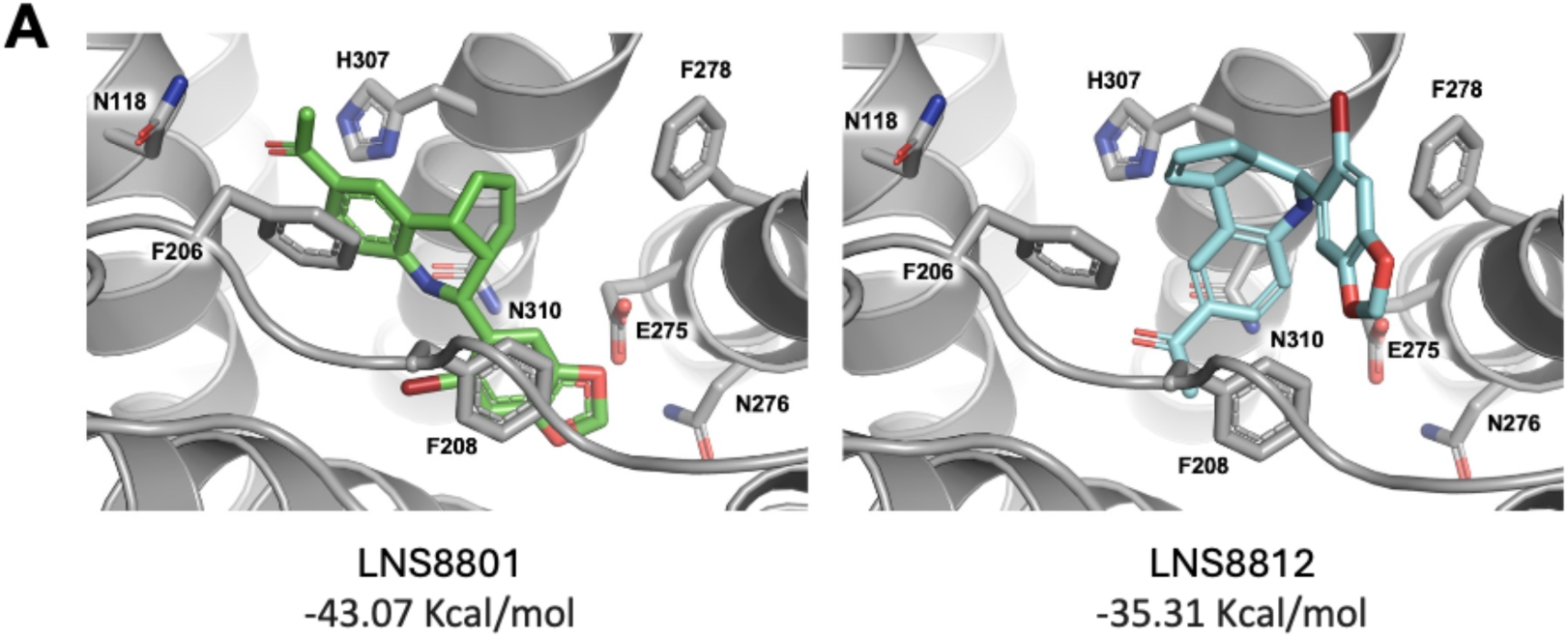
MM-GBSA modeling of LNS8801 and LNS8812 binding to GPER.

We evaluated both LNS8801 and LNS8812 using the Eurofins DiscoverX SAFETYscan off-target panel (**Supplemental Table 1**). LNS8801 had weak off-target activity on CNR1 (IC50 2.5 uM), HTR2A (IC50 8.2 uM), OPRM1 (IC50 3.5 uM), and OPRD1 (EC50 0.87 uM). The concentrations required for LNS8801 off-target binding far exceed the plasma levels required for efficacy in vivo (**Figure 2G**). LNS8812 had off-target activity on ADRA2A (IC50 2 uM), CNR1 (IC50 3 uM), HTR1A (IC50 2 uM), and AR (IC50 4.8 uM). While LNS8812 did not display any measurable activity in our cancer models (**Figure 1D-E**, **Figure 2A-B**), it may have biological activity in other settings.

### The activity of LNS8801 is dependent on GPER

To validate the on-target activity of LNS8801, CRISPR-Cas9 was used to deplete GPER in the YUMM1.7 and 2838c3 cancer lines. GPER protein depletion was confirmed using western blot and proliferation assays on GPER-depleted and non-depleted cells with LNS8801. Genetic depletion of GPER rendered YUMM1.7 and 2838c3 cells resistant to treatment, demonstrating the necessity of GPER for LNS8801 activity (**Figure 4A-B**). Consistent with this finding, GPER was also necessary for G-1 and LNS8801 cAMP induction as shown using siRNA knock-down (**Supplemental Figure 1B-C**).

**Figure 4:**
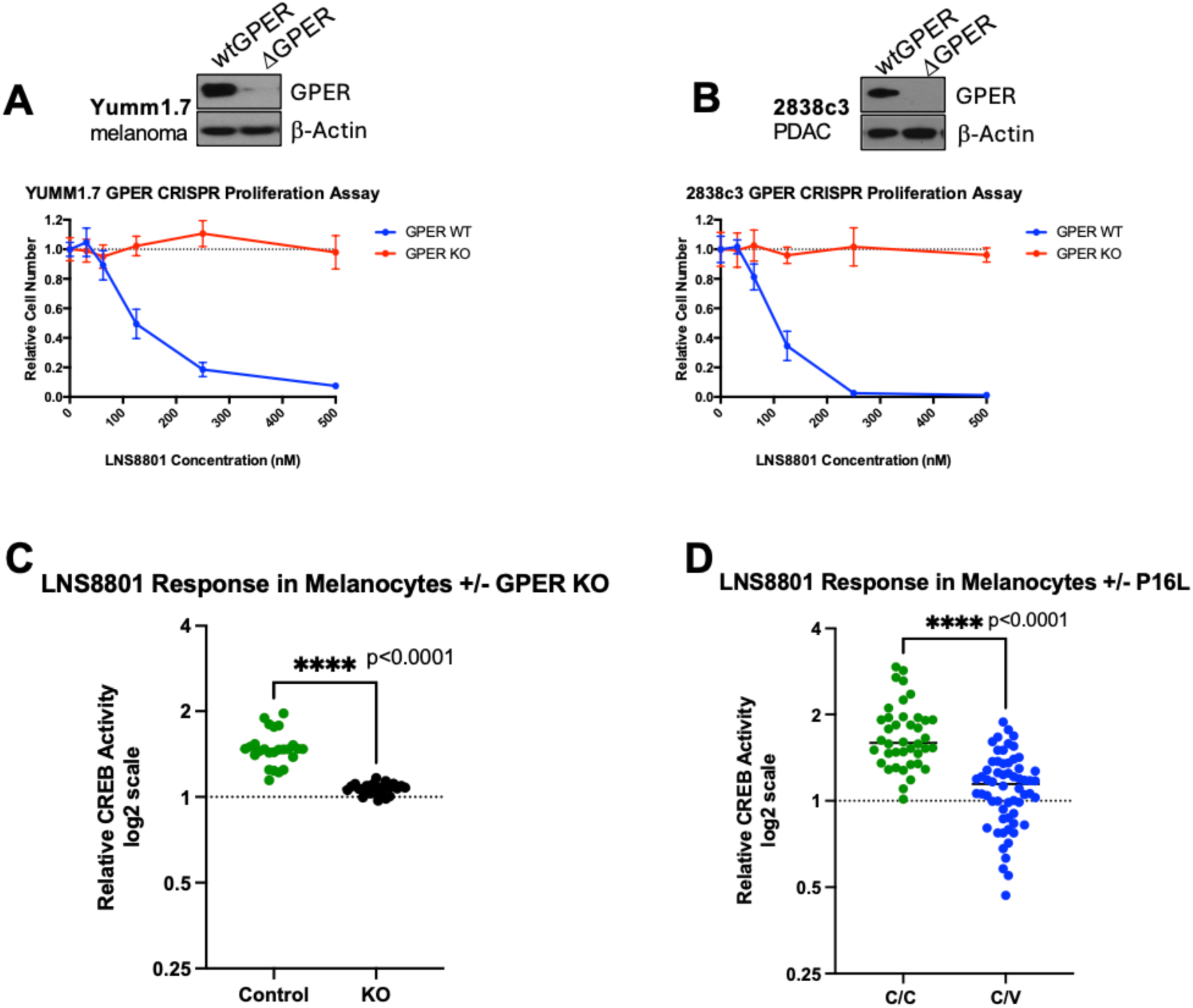
GPER is necessary, and genetic variation impacts LNS8801 effects. (A) Western blot demonstrating CRISPR-Cas9 GPER depletion and LNS8801 dose-response proliferation assay in YUMM 1.7 cells (melanoma). (B) Western blot demonstrating CRISPR-Cas9 GPER depletion and LNS8801 dose-response proliferation assay in 2838c3 cells (pancreatic cancer). (C) CREB reporter assay of human melanocytes with CRISPR-Cas9 depleted GPER treated with LNS8801, significance by the Mann-Whitney test. (D) CREB reporter assay of human melanocytes homozygous for consensus germline GPER (C/C) and heterozygous for variant germline GPER (C/V) treated with LNS8801, significance by the Mann-Whitney test.

These findings were further validated in primary human cells. GPER activation results in downstream activation of cAMP-response element-binding protein (CREB) in primary human melanocytes and other cell types (23,25). CRISPR-Cas9 was used to ablate GPER (**Supplemental Figure 2A-C**) and CREB activity was measured using a stably transduced CREB-luciferase reporter (26). This demonstrated that GPER was necessary for CREB signaling in response to LNS8801 (**Figure 4C**).

### Common germline variation impacts GPER activation with LNS8801

In humans, protein-coding, variant alleles of GPER are known. The most prevalent allele, rs11544331, is of particular importance as its frequency in White Europeans is estimated at 25%, such that roughly 50% of the population has at least 1 variant copy of GPER. The rs11544331 allele gives rise to single nucleotide polymorphism (SNP) involving replacement of the amino acid proline at position 16 in the N-terminus of GPER with leucine. Limited prior functional studies of this variant suggest that it is hypofunctional in response to G-1 (27,28). To further evaluate the effect of germline variations on LNS8801-induced CREB signaling, we determined the endogenous GPER sequence in a series of primary human melanocytes from single donors and identified lines that were homozygous for the consensus (C/C) and heterozygous (C/V) for the germline variation in GPER. We did not identify any lines homozygous for the variant, which is predicted to be in <5% of the genetically diverse population in Philadelphia where the donor skin tissues were obtained (27).

In total, 8 C/C and 12 C/V primary melanocyte lines were isolated. We treated these cells with LNS8801 and measured CREB activity using a stably transduced CREB reporter (26). LNS8801 treatment of GPER C/C melanocytes exhibited significantly stronger CREB induction than in GPER C/V melanocytes, indicating that the Pro16Leu variant attenuates GPER signaling in response to LNS8801 and may thereby render LNS8801 less efficacious in people (**Figure 3D**).

### GPER expression in normal tissues and select cancers using immunohistochemistry

Recently, a novel rabbit monoclonal antihuman GPER antibody, 20H15L21, was identified (29). The specificity of this antibody for GPER was determined using western blot analyses, immunofluorescence, and genetic manipulation of GPER. This antibody was used to determine GPER protein expression in 33 distinct normal tissues, with 3 cases for each tissue type (**Figure 5**). The normal tissue types with GPER protein expression in >50% of the cells were cerebrum, tonsil (epithelium), liver, small intestine (epithelium), kidney, pituitary gland, and colon (epithelium). The extent of GPER protein expression was also assessed in 6 different cancer types, with more than 30 cases for each cancer type (**Figure 6**). GPER protein was detectable in most colon cancers (81%), followed by pancreatic cancers (76%), melanomas (57%), and lung cancers (45%). In contrast, GPER expression was low in prostate cancers (6%) and not detected in 31 breast cancers.

**Figure 5:**
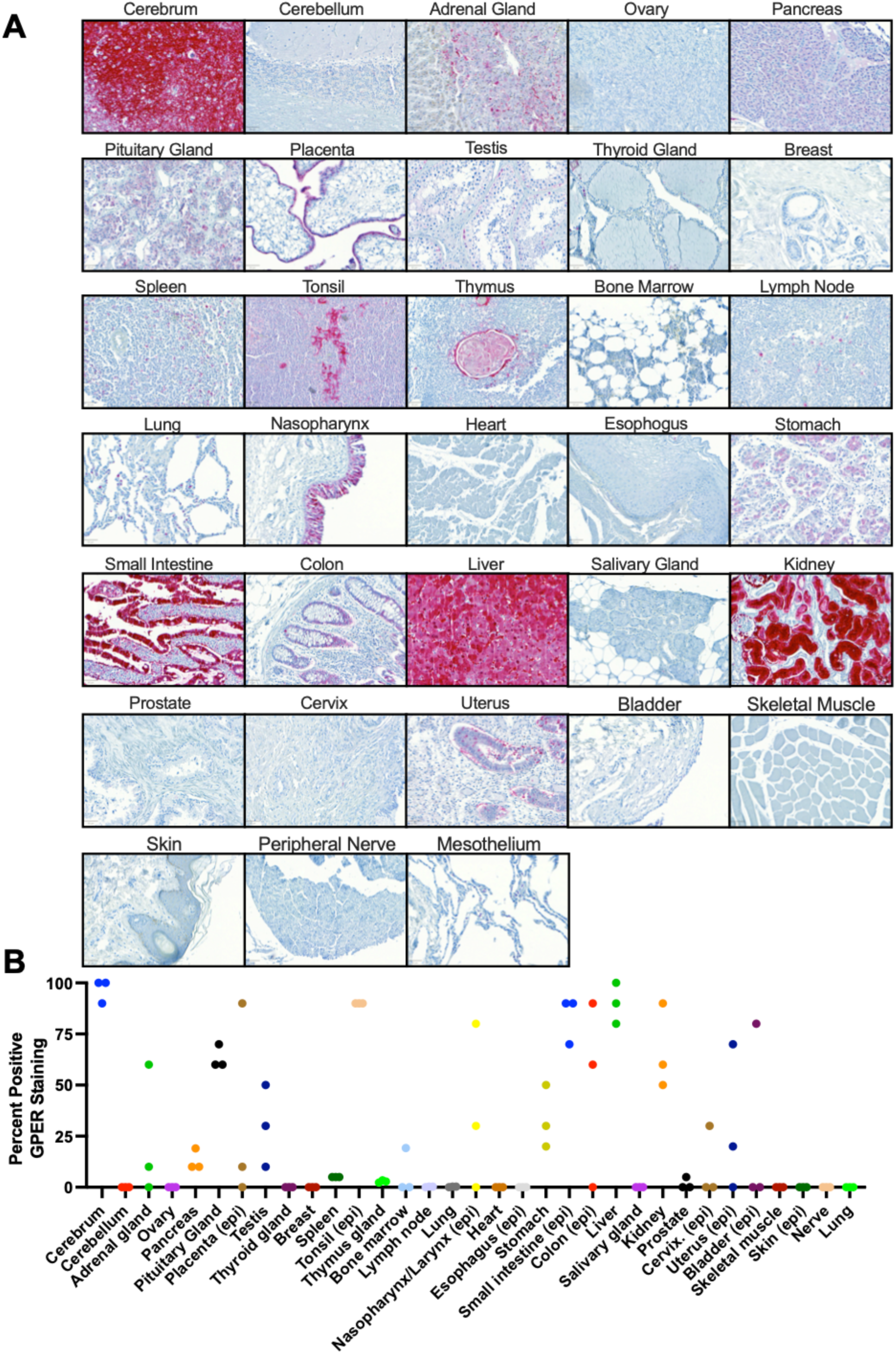
GPER staining in normal tissues. (A) Representative images of GPER-positive staining areas in normal tissues, scale bar = 50μm. (B) % of non-cancerous tissue samples with GPER staining across normal tissues; epithelial tissues were quantitated with staining in the epithelial region of the tissue specifically, and denoted with (epi).

**Figure 6:**
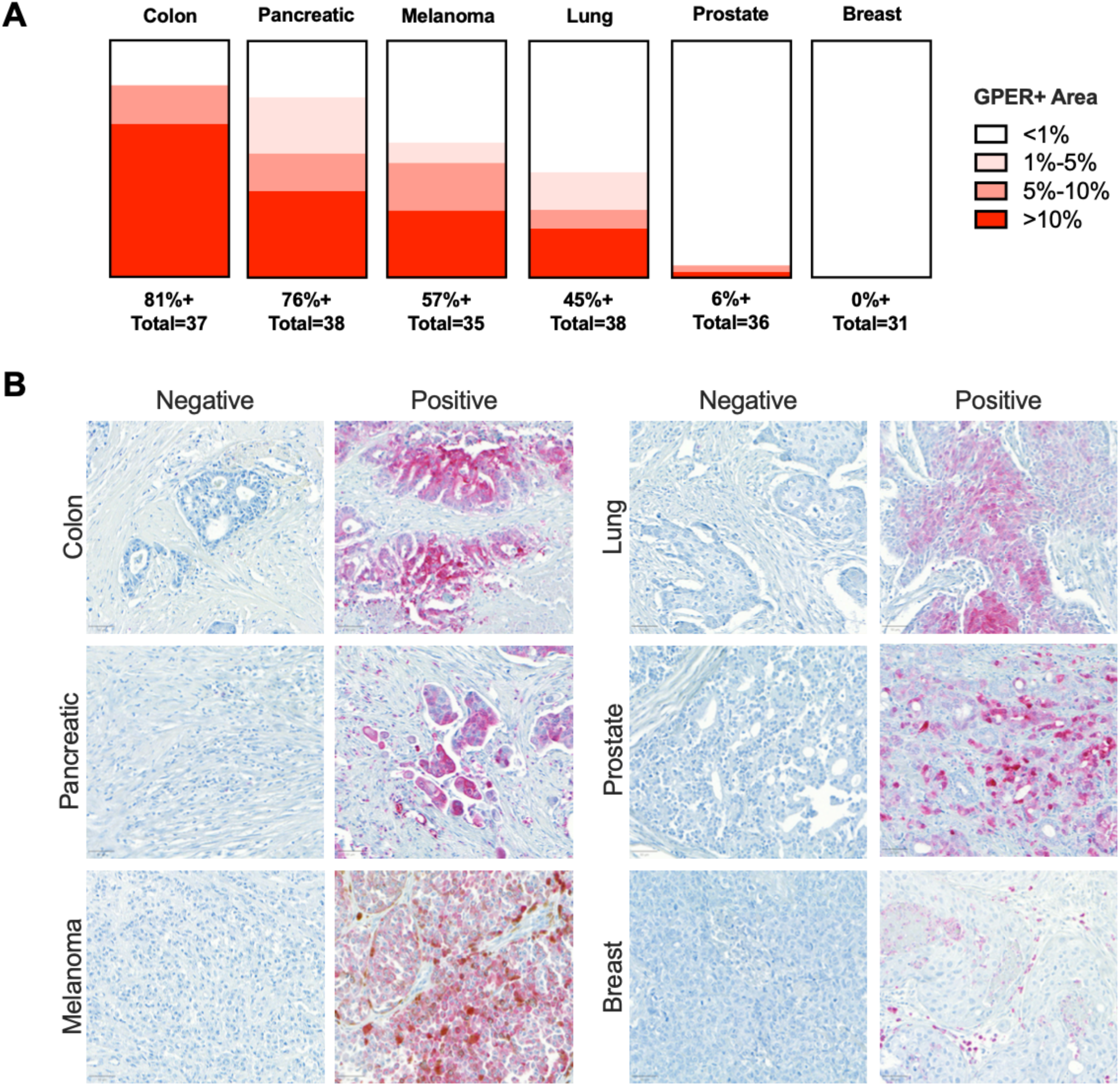
GPER staining in select malignancies. (A) Visualization of GPER staining across colon, pancreatic, melanoma, lung, prostate, and breast cancer cases. (B) Representative images of negative and positive GPER staining in the examined cancer types, scale bar = 50μm.

Although adjacent normal and cancer tissue could not be assessed in these samples, there appears to be a concordance of GPER protein expression between normal and cancer tissues. For example, normal colon and pancreas tissue expresses GPER protein, and a high percentage of colon and pancreatic cancer cases also expresses GPER protein. Conversely, we observed low GPER protein expression in both normal breast and prostate and a similarly low GPER protein expression in breast and prostate cancers.

## Discussion

The racemic GPER agonist G-1 has been an invaluable tool for studying GPER (15) because it does not simultaneously bind to or activate classical nuclear estrogen receptors. However, G-1 is racemic, and the question of what the activity of each enantiomer of G-1 is has never been addressed. Through the isolation of pure samples of each enantiomer followed by in vitro and in vivo testing, we have shown that the desirable biological activity results from a single enantiomer. Through single crystal x-ray structural analysis, we have unambiguously determined the configuration of the active enantiomer, named LNS8801, as 1-[(3aS,4R,9bR)-4-(6-bromo-1,3-benzodioxol-5-yl)-3a,4,5,9b-tetrahydro-3H-cyclopenta[c]quinolin-8-yl] ethanone. Subsequently, we have demonstrated that LNS8801 is orally bioavailable and has potent antitumor activity in vivo at low doses. Three independent approaches demonstrated that LNS8801 signaling and activity require GPER protein expression.

Molecular modeling of the binding of each of the enantiomers of racemic G-1 with GPER correctly predicts that the LNS8801 enantiomer should exhibit much greater activity than the LNS8812 enantiomer, consistent with the experimentally observed results. The original screening study resulting in the identification of racemic G-1 as well as subsequent synthetic work have resulted in a variety of structural G-1 analogs (15,22,30). In all these studies, the synthetic approach resulted in the isolation of the materials as racemic compounds. In testing, some of these compounds behave as agonists of GPER, while others behave as antagonists. Given our results that show that the pure enantiomers of racemic G-1 display very different activities, it is very likely that the pure enantiomers of each of these racemic analogs of G-1 will also exhibit different activities with respect to GPER and that molecular modeling may provide a useful way of predicting the activity of specific enantiomers.

The fact that G-1 is racemic impacts the interpretation of prior published literature that used G-1 to assess functional consequences of selective GPER activation. While much of the literature is consistent (1), there have been some contradictory studies related to the impact of GPER activation. Although the inactive enantiomer LNS8812 did not demonstrate measurable effects in the cancer models used in this report (**Figures 1D-E 2A-B**), it did demonstrate differences in the off-target binding screen (**Supplemental Table 1**). Compared with LNS8801, LNS8812 had distinct inhibitory activity on ADRA2A, HTR1A, and the androgen receptor and possibly other untested biological mechanisms. Although our studies were not designed to assess this directly, it is possible that LNS8812 has activity in certain biological contexts, and thus racemic G-1 may have activities that are unrelated to GPER. For example, LNS8801 demonstrated significantly improved activity in vivo compared with G-1 (**Figure 2 A-B)** at doses well above saturation (**Figure 2 E-F)**, suggesting a greater than 2-fold increase in activity. Additionally, recent studies have demonstrated more consistent beneficial activity of LNS8801 compared with G-1 on high blood pressure (31). Because we have demonstrated that the biologically significant activity of racemic G-1 resides in a single enantiomer LNS8801 and that the use of racemic G-1 in studies seeking to eluate the activity of GPER can be complicated by the presence of the inactive enantiomer LNS8812, we recommend caution when interpreting results of experiments using G-1. In addition, complementary approaches using GPER protein depletion should be performed to confirm the on-target activity of the GPER agonist effects.

We described GPER protein expression in 33 normal tissue types and 6 common cancer types. The antibody labeling was robust in the IHC application, demonstrating a very low background and a wide dynamic range, labeling GPER in both the cytoplasm and the plasma membrane. Our findings are consistent with and validate the initial report of the monoclonal antihuman GPER antibody, clone 20H15L21 (29). These data extend the findings from that prior report and show that GPER protein is expressed in a high percentage of colon, pancreatic, melanoma, and lung cancer cases, consistent with reports showing efficacy in these preclinical models (3,4,17–19). While GPER expression was detected in a high percentage of these cancer types, we observed heterogeneity between cases of the same cancer type, suggesting that GPER protein expression in patient samples may be a useful biomarker for GPER-directed therapies (32,33).

Additionally, both studies detected little to no GPER in prostate and breast cancers. This is noteworthy as there are older reports suggesting a potential role for GPER in preclinical models of breast cancer (34). Although GPER signaling was originally reported to be tumor promoting in some breast cancer models (35), subsequent reports show that GPER signaling inhibits breast cancer (36–38). Additionally, several studies have documented correlations between GPER expression and prognosis in breast cancer, with contradictory results (39,40). Critically, those studies did not determine the amount of GPER signaling in the studied tumors and whether the patients harbored the germline GPER variant. It is therefore still unclear whether GPER plays a significant role in spontaneous human breast cancer. Given the wide range of GPER-mediated functions that have been identified in various tissue types, including vascular elasticity and blood pressure, lipid and glucose metabolism, and inflammation (1), assays and reagents that can reliably detect and selectively modulate GPER activity in human tissues are critically important to further the understanding of GPER biology.

A critical aspect to consider moving forward is the potential functional impact of GPER variants, particularly the Pro16Leu variant, which appears to be unique to humans and absent in rodents and primates. The allele frequency is 25% in White Europeans, such that roughly 50% of that population has at least 1 variant copy of GPER. Historically, germline GPER variants were not known to impact GPER function; however, studies associating GPER variations with higher blood pressure (27) suggest that common germline variations may influence GPER activation and signaling. The impact of variant GPER genes is largely unexplored, though 1 prior report suggests that GPER P16L is hypofunctional (27), consistent with our findings here using primary human cells expressing the endogenous GPER genes and the highly specific GPER agonist, LNS8801.

This study has a significant translational impact on the development of therapies targeting GPER. Since the desirable biological activity of racemic G-1 resides in only 1 enantiomer, LNS8801, Linnaeus Therapeutics has been pursuing clinical development of this pure enantiomer. Thus, LNS8801 has been evaluated in a multisite phase 1 clinical trial (NCT04130516) and has demonstrated benefit in patients with multiple cancer types, including melanoma and lung and colon cancer, among others (32,33,41–43). Early clinical data from the LNS8801 trial indicate that patients with the consensus germline GPER genotype are associated with improved overall survival compared with patients with 1 or 2 variant alleles (41–43). If validated in expanded clinical trials, GPER sequence and expression may prove to be predictive biomarkers that prospectively identify patients most likely to benefit from LNS8801 therapy.

## Acknowledgments

The authors would like to thank Patrick Mooney for their critical presubmission review of this manuscript. The authors would also like to thank Bryan Szpunar for overseeing this project at DCL Pathology.

## Methods

### Cells, Tissue Culture, and Reagents

YUMM1.7 cells were purchased from ATCC (CRL-3362); 2838c3 cells were a gift from female mice in the laboratory of Ben Stanger (University of Pennsylvania); HL-60 cells were purchased from ATCC (CCL-240); and HEK293T cells were purchased from ATCC (CRL-3216). 2838c3, YUMM1.7, and 293T cells were cultured in DMEM medium (ATCC 30-2002) supplemented with 5% fetal bovine serum (Cat # MT35-010-CV), 100 units/mL penicillin, and 100 μg/mL streptomycin (Cat # 15140122). HL-60 cells were cultured in RPMI 1640 medium (Gibco, Cat# 61870036) supplemented with 10% fetal bovine serum (Gibco, Cat# A5670701), 100 units/mL penicillin, and 100 μg/mL streptomycin (Gibco, Cat# 15140122). Primary melanocytes were derived samples from freshly discarded human foreskin obtained through the Skin Translational Research Core within the Skin Biology and Diseases Resource-based Center (SBDRC) at Penn Medicine. After overnight incubation in Dispase (Cat # 354235), the epidermis was mechanically separated from the dermis. The epidermal layer was then enzymatically dissociated in 0.25% Trypsin-EDTA (Cat # 25200056) at 37°C for 8 minutes, followed by 2 minutes of vigorous shaking. Trypsin activity was quenched with DMEM 5%. Cells were pelleted and plated in Media 254 (Cat # M254500) supplemented with Human Melanocyte Growth Supplement (Cat # S0025) as well as 100 units/mL penicillin and 100 μg/mL streptomycin (Cat # 15140122). All cells were maintained at 37°C in 5% CO_2_.

LNS8801 and LNS8812 were provided by Linnaeus Therapeutics. G-1 was isolated using chiral chromatography, resulting in LNS8801 and LNS8812. IBMX, G-1, and α-MSH were purchased from Cayman Chemical Company. D-luciferin potassium salt was purchased from Gold Biotechnology.

### Mice

All mice were purchased from Jackson Laboratories (Bar Harbor, ME). Five-to 7-week-old C57BL/6J or nude (NU/J) mice were allowed to acclimatize for 1 week before being used for experiments. All mice were female unless otherwise noted. These studies were performed without inclusion/exclusion criteria or blinding but included randomization. Based on a 2-fold anticipated effect, we performed experiments with at least 5 biological replicates. All procedures were performed in accordance with International Animal Care and Use Committee–approved protocols at the University of Pennsylvania.

### Subcutaneous Tumors and Treatments

Subcutaneous tumors were initiated by injecting tumor cells in 50% Matrigel (Corning, Bedford, MA) into the subcutaneous space on the left and right flanks of mice, and 2 × 10^5^ 2838c3 murine PDAC cells were used for each tumor. Subcutaneous treatments were prepared by first dissolving compounds in 100% ethanol at a concentration of 1 mg/mL. The desired amount of compound for 5 injections, given the dose, was then mixed with a 250-**μ**L volume of sesame oil, and the ethanol was evaporated off using a Savant Speed Vac (Thermo Fisher Scientific, Waltham, MA, USA). G-1 injections of 50-**μ**L were delivered into the subcutaneous space using a 28G x 1/2 inch insulin syringe on the dorsal region of the mouse. Soluble oral treatments were prepared by dissolving compound in 10% DMSO and 90% PEG400, and 100-**μ**L volumes were used for oral delivery. Insoluble oral treatments were prepared by suspending compound in 0.5% hydroxypropyl-methyl cellulose in water, and 100-**μ**L volumes were used for oral treatment. Oral treatments were delivered by gavage. Oral oil formulations were prepared by diluting LNS8801 in DMSO, resulting in a 10-mg/mL solution. A fresh 10-mg/mL DMSO stock was prepared at the beginning of each treatment pulse. This LNS8801 solution was used to prepare 0.01-mg/kg, 0.1-mg/kg, and 1-mg/kg oral doses by mixing with additional DMSO, followed by EtOH and sesame oil. 100 **μ**L of these solutions were delivered by oral gavage. The final vehicle consisted of 13% DMSO, 5% EtOH, and 82% sesame oil. Treatments were delivered for 3 consecutive days, over 3 consecutive weeks, resulting in 9 doses total.

### Survival Analysis

As subcutaneous tumors grew in mice, perpendicular tumor diameters were measured using calipers. Volume was calculated using the formula L × Ŵ2 × 0.52, where L is the longest dimension and W is the perpendicular dimension. Animals were euthanized when tumors exceeded a protocol-specified size of 15 mm in the longest dimension. Secondary endpoints included severe ulceration, death, and any other condition that falls within the International Animal Care and Use Committee Guidelines for Rodent Tumor and Cancer Models at the University of Pennsylvania.

### Predictive Modeling

The predictive binding energies for LNS8801 and LNS8812 on GPER were calculated using molecular mechanics energies combined with generalized Born and surface area continuum solvation using Schrodinger Maestro (MM-GBSA) and using methodology as previously described (44). The absolute values calculated are not necessarily in agreement with experimental binding affinities. However, the ranking of the ligands based on the calculated binding energies (MMGBSA DG Bind) can be expected to agree reasonably well with the ranking based on experimental binding affinity, particularly in the case of congeneric series. As the MM-GBSA binding energies are approximate free energies of binding, a more negative value indicates stronger binding.

### Pharmacokinetic Analysis

Pharmacokinetic analysis was performed by WuXi AppTec (Cranbury, NJ), Briefly, 5 male and 5 female C57BL/6 mice were treated with LNS8801 at 0.1 mg/kg via oral gavage once daily for 3 consecutive days. LNS8801 was formulated in 13% DMSO, 5% ethanol, and 82% sesame oil at the concentration of 0.026 mg/mL and administered at 3.8 mL/kg. Blood samples were collected on study day 1 and day 3 post dosing at 0.25, 0.5, 1, 2, 4, 8, and 24 hours, respectively. An extra blood sample was taken on study day 3 immediately prior to dosing. Concentrations of LNS8801 in the plasma were determined by LC-MS/MS. The bioanalytical assay for LNS8801 provided an LLOQ of 0.030 ng/mL and a linear range of up to 100 ng/mL. A noncompartmental model was used to calculate pharmacokinetic parameters using Phoenix WinNonlin 6.3.

### Transfections and Transduction

HEK293T cells were seeded to 60% confluency in 6-well plates in 2 mL of DMEM 5%. For each well, 1.22 µg of the lentiviral vector was mixed with viral packaging plasmids pCMVΔR8.91 (0.915 µg) and pUC-MDG (0.305 µg). The plasmid solution, along with 7.32 µL of Promega’s FuGENE® 4K Transfection Reagent (3:1 ratio of FuGENE to DNA), was added to serum-free DMEM for a final volume of 100 µL. The resulting mixture was incubated at room temperature for 15 minutes and then added dropwise to the HEK293T culture media.

Twenty-four hours after transfection, the culture media was replaced with 4 mL of DMEM 5%. At 48 hours, viral supernatant was collected and filtered through a 0.45-µm syringe filter with polyethersulfone (PES) membrane (MilliporeSigma). Melanocytes seeded at 1.5 × 10^5^ cells per well were incubated in the viral supernatant in the presence of 5 µg mL^-1^ polybrene. The cells were centrifuged at 300 g for 60 minutes at room temperature, followed by 25 minutes of incubation at 37°C. The viral supernatant was then removed and replaced with fresh melanocyte growth media.

### Genetic Depletions

Murine guide RNAs were designed using the CHOPCHOP CRISPR design tool (https://chopchop.cbu.uib.no/). Five guide RNA sequences for GPER knockout were screened, with 1 guide sequence being the most effective and used for further experiments:

5’-GATGCCCCGGGGAACCTCAC-3’

We used lentiviral transduction to deliver dox-inducible Cas9 (45) and gRNA targeting GPER of murine YUMM1.7 melanoma and 2838c3 PDAC cells. Transduced cells were selected with puromycin, and single cells were subsequently isolated, expanded, and examined for GPER protein expression, compared with clones isolated in parallel with no doxycycline treatment.

Human guide RNAs were designed using VectorBuilder’s Mammalian CRISPR Lentiviral Vector System. The guide RNA sequences for GPER knockout are:

Guide 1: 5′-GCAGTACGTGATCGGCCTGT-3′

Guide 2: 5′-GTTCCGCACCAAGCACCACG-3′

For the control group, Scramble guides targeting nongenomic or safe-harbor regions were used:

Nongenomic guide: 5’-GTGTAGTTCGACCATTCGTG-3’

AAVS1 safe-harbor guide: 5’-GGGGCCACTAGGGACAGGAT-3’

Equal amounts (660 ng) of each GPER knockout guide RNA were combined into the viral transfection mix to generate an ∼350 base pair deletion in the GPER coding region of the cells to be infected. LentiCRISPR transductions in human melanocytes were conducted as previously described for the other lentiviral constructs used in this study. Successful GPER knockout was validated via PCR amplification of the GPER gene from genomic DNA followed by Sanger Sequencing (Supp. Fig 2A-C).

GPER small interfering RNA (siRNA) was purchased from Santa Cruz Biotechnology (Dallas, TX), sc-60743. HL-60 cells were centrifuged and plated in a 6-well plate at 400,000 cells/well. Transfection of the siRNA was performed following the protocol of X-tremeGene siRNA transfection reagent from Roche Diagnostics GmbH (Mannheim, Germany) and Opti-MEM. The transfection reagent and GPER siRNA (500 pmol and 1000 pmol) were incubated at room temperature for 20 minutes and then added to 2 mL of cell suspension in wells. The transfected cells were then incubated at 37°C under 5% CO_2_ for 48 hours. HL-60 control cells were treated with the transfection reagent with or without a non-GPER siRNA. After 48 hours, cells were used for analysis.

### Sanger Sequencing

Genomic DNA was isolated using the Qiagen DNeasy Blood & Tissue Kit. A PCR reaction was carried out using 20 ng of genomic DNA, Q5^®^ Hot Start High-Fidelity DNA Polymerase (Cat # M0493S), GeneAmp™ 10 mM dNTP Blend (Cat # N8080260), Q5^®^ High GC Enhancer (Cat # B9028AVIAL), Q5^®^ Reaction Buffer (Cat # B9027SVIAL), and a final concentration of 500 nM of 2 self-designed primers that amplify a section of GPER’s coding region. The GPER PCR primer sequences are as follows:

GPER forward primer: 5’-TTCCTGTCTGACAAATGCCAGG-3’

GPER reverse primer: 5’-TGATGAAGACGTTCTCCGGC-3’

The PCR reaction was run out on a 1% agarose gel and visualized on the BioRad ChemiDoc Imaging System. The target band was excised and sent for Sanger Sequencing, which was performed by the DNA Sequencing Facility at Penn Medicine. The sequencing data files indicated successful GPER knockout in cells infected with the GPER knockout guides (Supp. Fig 2C) compared with cells infected with the Scramble guide (Supp. Fig 2B).

### cAMP Assay

A homogeneous time-resolved fluorescence (HTRF) for cAMP from CisBio (Bedford, MA) was used following the manufacturer’s suggested protocol. HL-60 cells were grown in 10% charcoal-stripped FBS RPMI media for 72 hours prior to assays. Cells were centrifuged and counted with a hemocytometer from Hausser Scientific (Horsham, PA). The total number of cells needed to complete the assay was determined based on 8,000 cells/well. Cells were diluted in a 5:1 (5X) stimulation buffer containing 500 µM of 3-isobutyl-1-methylxanthine (IBMX) (Sigma Aldrich, Saint Louis, MO). To an HTRF 96-well low volume white plate (CisBio), 2 µL of (5X) agonist was added to the designated wells, followed by 8 µL of cold cell suspension. Two controls were performed during the experiment to establish basal cAMP formation and maximum cAMP. The maximum level of cAMP was determined with forskolin (20 µM). Cells were covered with a clear plastic film and incubated at 37°C for 15 mins. During the 15-minute incubation, a 4:1 solution of lysis buffer to cryptate-tagged cAMP or d2 tagged monoclonal antibody was prepared individually. After the 15-minute incubation period, 5 µL of lysis buffer/cAMP-d2 (acceptor) and 5 µL of lysis buffer/monoclonal anti-cAMP Eu3+ cryptate (donor) were added to all wells. Once added, the plate was sealed, covered with aluminum foil, and incubated at room temperature for 30 minutes. After the allotted time, cells were read using Flexstation3 and with ex 314 nm and em 665 nm/em 620 nm, with an integration delay of 50 µs and an integration of 400 µs. The signal from agonism was normalized to the forskolin control. Nonlinear regression curves were then generated using Prism (GraphPad, La Jolla, CA) to determine EC50 values. A triplicate was performed to obtain an n=3. Standard deviations amongst the calculated EC50 values were calculated using Prism.

### CREB-Luciferase Assays

A CREB-luciferase assay is a reliable in vitro biomolecular assay that uses a luciferase reporter gene to monitor the activity of the cyclic-AMP response element-binding protein (CREB) (46). After transducing melanocytes with the CREB-luciferase reporter, they were plated in white, clear bottom 96-well plates (Cat # 62-1076-1) at a density of 18,000 cells per well and treated with ligands overnight in 180 µL of Media 254 with dialyzed serum. Serum was dialyzed prior to adding to Media 254 using the 3,500 MWCO Slide-A-Lyzer™ Dialysis Cassette (Cat # 66333) to remove MSH, PMA, and other small molecules in the serum that elevated basal CREB activity in these cells. IBMX was used in all conditions at a concentration of 100 µM to prevent cAMP degradation. α-MSH was used as a positive control. About 18 hours after ligand treatment, a final concentration of 150 µg/mL of D-luciferin potassium salt was added to the wells, and luminescence was measured using the BioTek Synergy H1 Multimode Reader.

### Tissue Microarrays

FFPE tissue microarrays were purchased from Biomax (Derwood, MD, USA). Catalog numbers for the tissue arrays are as follows: Normal tissue (FDA999y2), melanoma (ME1002b), and multiple other cancers (BC000119b).

### GPER Immunohistochemistry and Imaging

Tissue microarrays were stained using the GPER recombinant rabbit monoclonal antibody clone:20H15L21 (Thermo Fisher Scientific, catalog #703480) by DCL Pathology (Indianapolis, IN, USA). IHC staining for GPER was performed using the Ventana Benchmark Ultra platform (Roche Diagnostics). Tissue sections were cut onto positively charged slides at a thickness of 3 to 5 microns and baked at 60°C to 65°C for 10 minutes prior to IHC staining. Slides were deparaffinized on the Benchmark Ultra at 72°C using EZ Prep solution (Cat. No. 950-102). Slides were then treated with Ultra Cell Conditioning Solution 1 (950–224) for 36 minutes at 95°C. The antibody used for this assay was anti-GPER (20H15L21) rabbit monoclonal antibody and was diluted 1:375 in Ventana Antibody Diluent (Cat. No. 251-018). 100 μL of diluted antibody was applied to the slides and incubated for 32 minutes at 37°C. After the primary antibody incubation, the slides were rinsed, and detection was performed using the Roche ultraView Universal Alkaline Phosphatase Red Detection Kit (Cat. No. 760-501). After the detection reaction was completed, tissues were counterstained with Hematoxylin II (Cat. No. 790-2008) for 4 minutes and blued with Bluing Reagent (Cat. No. 760-2037) for 4 minutes. Slides were then rinsed with soapy water followed by DI water and dehydrated in 100% reagent alcohol. Slides were transferred to xylene and coverslipped. Stained slides were Whole Slide Imaged using the Motic EasyScanPro platform and stored as Leica Aperio SVS files.

### Immunohistochemistry Quantification

Scanned slide images were viewed with QuPath, (47) and images of the entire core were exported for all evaluable, nondefective cores. Core images were then assessed for the extent of GPER staining by calculating the area of GPER staining using Adobe Photoshop. The total area of the core was measured in pixels, subtracting any acellular regions representing >10% of the core area. The area of GPER staining was measured in pixels using a color range that captures red staining indicative of GPER protein expression. The area of GPER staining was divided by the cellularized area of the tissue core and multiplied by 100, resulting in a percent GPER staining metric. Tissue-staining results were reviewed by a board-certified pathologist and adjusted when necessary to account for staining in the normal or cancer tissue of interest. All quantification data are in Supplementary Data Table 1.

### Statistical Analysis

All statistical analysis was performed using GraphPad Prism 10. The details of each statistical test used are included in the figure legends.

---

**Supplemental Figure 1:**
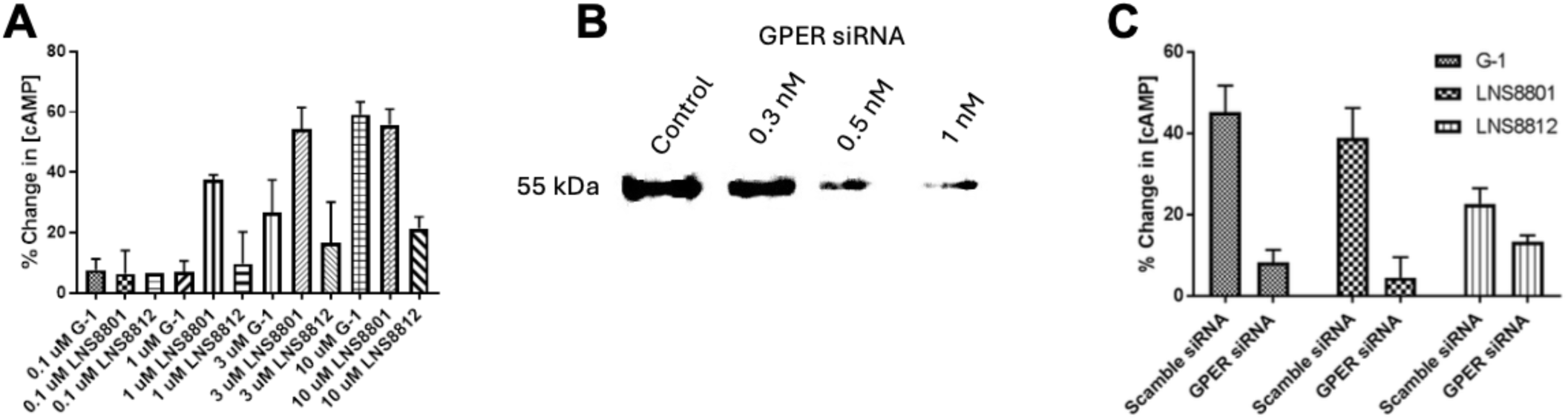
Validation of LNS8801 activity and necessity of GPER in HL-60 cells. (A) Dose response of cAMP assay in HL-60 cells treated with G-1, LNS8801, and LNS8812 relative to control. (B) Western blot demonstrating GPER depletion by anti-GPER siRNA in HL-60 cells. (C) cAMP assay of HL-60 cells with and without GPER siRNA depletion treated with G-1, LNS8801, and LNS8812.

**Supplemental Figure 2:**
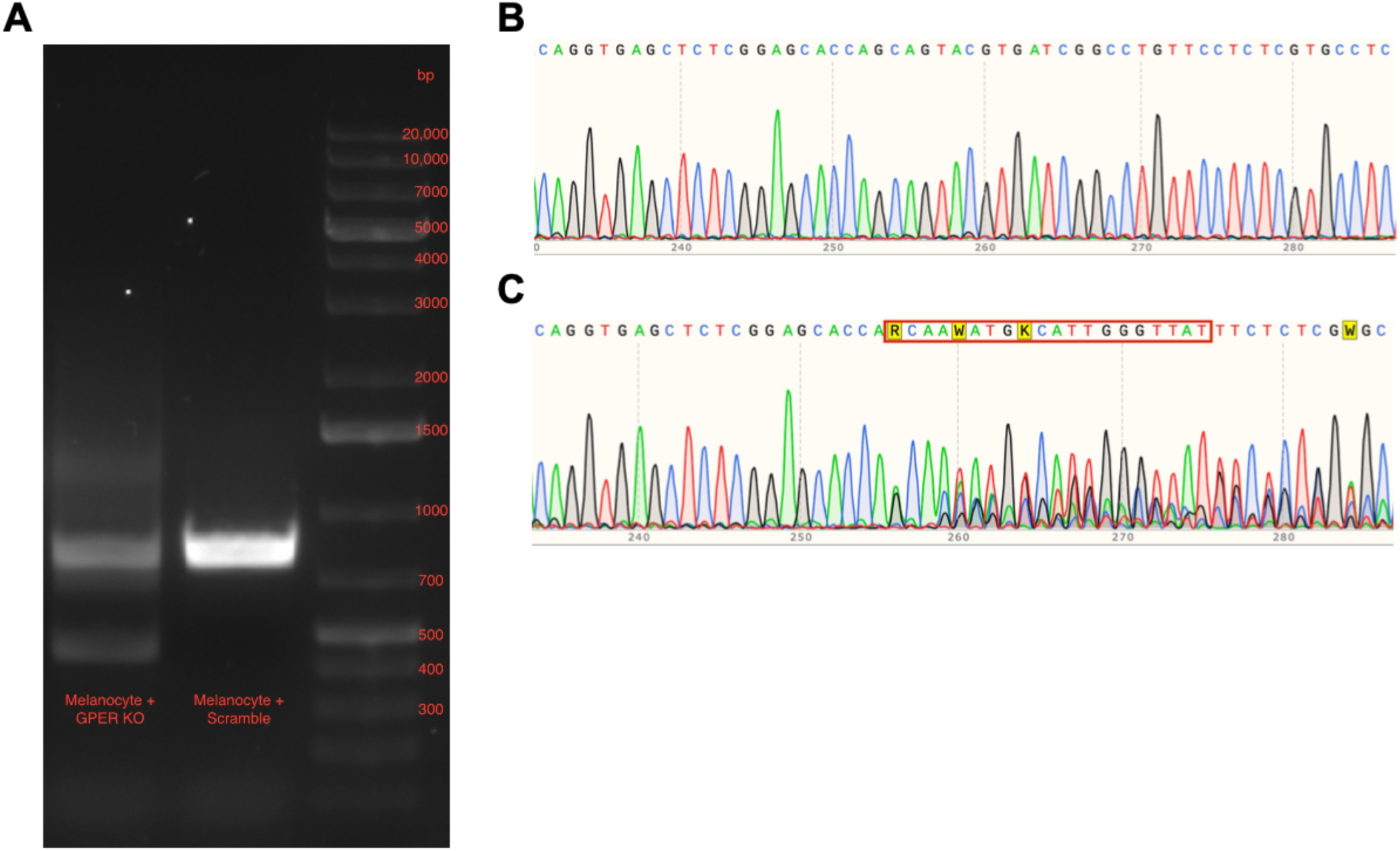
CRISPR-Cas9 depletion of GPER in human melanocytes. (A) Agarose gel of PCR-amplified GPER from genomic DNA in control or GPER-depleted cells via CRISPR-Cas9. (B) Sanger sequencing of cells targeted with control CRISPR-Cas9. (C) Sanger sequencing of cells targeted with GPER CRISPR-Cas9.

**Supplemental Table 1:** Off-target binding of LNS8801 and LNS8812 using Eurofins Discover X.

| Compound Name | Order ID | Target Class | Assay Name | Mode | Assay Target | Result Type | Value Prefix | RC50 (μM) | Hill | Curve Bottom | Curve Top | Max Response |
| --- | --- | --- | --- | --- | --- | --- | --- | --- | --- | --- | --- | --- |
| LNS8801 | US073-0000619-O | GPCR | Calcium Flux | Agonist | ADORA2A | EC50 | > | 10 |  |  |  | 15.02 |
| LNS8801 | US073-0000619-O | GPCR | Calcium Flux | Antagonist | ADORA2A | IC50 | > | 10 |  |  |  | 0 |
| LNS8801 | US073-0000619-O | GPCR | Calcium Flux | Agonist | ADRA1A | EC50 | > | 10 |  |  |  | 0.43 |
| LNS8801 | US073-0000619-O | GPCR | Calcium Flux | Antagonist | ADRA1A | IC50 | > | 10 |  |  |  | 0 |
| LNS8801 | US073-0000619-O | GPCR | cAMP | Agonist | ADRA2A | EC50 | > | 10 |  |  |  | 0 |
| LNS8801 | US073-0000619-O | GPCR | cAMP | Antagonist | ADRA2A | IC50 | > | 10 |  |  |  | 38.4 |
| LNS8801 | US073-0000619-O | GPCR | cAMP | Agonist | ADRB1 | EC50 | > | 10 |  |  |  | 3.08 |
| LNS8801 | US073-0000619-O | GPCR | cAMP | Antagonist | ADRB1 | IC50 | > | 10 |  |  |  | 34 |
| LNS8801 | US073-0000619-O | GPCR | cAMP | Agonist | ADRB2 | EC50 | > | 10 |  |  |  | 1.24 |
| LNS8801 | US073-0000619-O | GPCR | cAMP | Antagonist | ADRB2 | IC50 | > | 10 |  |  |  | 47.12 |
| LNS8801 | US073-0000619-O | GPCR | Calcium Flux | Agonist | AVPR1A | EC50 | > | 10 |  |  |  | 0 |
| LNS8801 | US073-0000619-O | GPCR | Calcium Flux | Antagonist | AVPR1A | IC50 | > | 10 |  |  |  | 30.41 |
| LNS8801 | US073-0000619-O | GPCR | Calcium Flux | Agonist | CCKAR | EC50 | > | 10 |  |  |  | 4.81 |
| LNS8801 | US073-0000619-O | GPCR | Calcium Flux | Antagonist | CCKAR | IC50 | > | 10 |  |  |  | 0 |
| LNS8801 | US073-0000619-O | GPCR | Calcium Flux | Agonist | CHRM1 | EC50 | > | 10 |  |  |  | 0 |
| LNS8801 | US073-0000619-O | GPCR | Calcium Flux | Antagonist | CHRM1 | IC50 | > | 10 |  |  |  | 32.48 |
| LNS8801 | US073-0000619-O | GPCR | cAMP | Agonist | CHRM2 | EC50 | > | 10 |  |  |  | 0 |
| LNS8801 | US073-0000619-O | GPCR | cAMP | Antagonist | CHRM2 | IC50 | > | 10 |  |  |  | 31.29 |
| LNS8801 | US073-0000619-O | GPCR | Calcium Flux | Agonist | CHRM3 | EC50 | > | 10 |  |  |  | 2.91 |
| LNS8801 | US073-0000619-O | GPCR | Calcium Flux | Antagonist | CHRM3 | IC50 | > | 10 |  |  |  | 4.12 |
| LNS8801 | US073-0000619-O | GPCR | cAMP | Agonist | CNR1 | EC50 | > | 10 |  |  |  | 0 |
| LNS8801 | US073-0000619-O | GPCR | cAMP | Antagonist | CNR1 | IC50 | = | 2.51137 | 1.56 | 0.22 | 55.62 | 49.99 |
| LNS8801 | US073-0000619-O | GPCR | cAMP | Agonist | CNR2 | EC50 | > | 10 |  |  |  | 0 |
| LNS8801 | US073-0000619-O | GPCR | cAMP | Antagonist | CNR2 | IC50 | > | 10 |  |  |  | 52.22 |
| LNS8801 | US073-0000619-O | GPCR | cAMP | Agonist | DRD1 | EC50 | > | 10 |  |  |  | 2.92 |
| LNS8801 | US073-0000619-O | GPCR | cAMP | Antagonist | DRD1 | IC50 | > | 10 |  |  |  | 23.72 |
| LNS8801 | US073-0000619-O | GPCR | cAMP | Agonist | DRD2S | EC50 | > | 10 |  |  |  | 2.53 |
| LNS8801 | US073-0000619-O | GPCR | cAMP | Antagonist | DRD2S | IC50 | > | 10 |  |  |  | 11.8 |
| LNS8801 | US073-0000619-O | GPCR | Calcium Flux | Agonist | EDNRA | EC50 | > | 10 |  |  |  | 0 |
| LNS8801 | US073-0000619-O | GPCR | Calcium Flux | Antagonist | EDNRA | IC50 | > | 10 |  |  |  | 4.94 |
| LNS8801 | US073-0000619-O | GPCR | Calcium Flux | Agonist | HRH1 | EC50 | > | 10 |  |  |  | 1.05 |
| LNS8801 | US073-0000619-O | GPCR | Calcium Flux | Antagonist | HRH1 | IC50 | > | 10 |  |  |  | 5.27 |
| LNS8801 | US073-0000619-O | GPCR | cAMP | Agonist | HRH2 | EC50 | > | 10 |  |  |  | 3.9 |
| LNS8801 | US073-0000619-O | GPCR | cAMP | Antagonist | HRH2 | IC50 | > | 10 |  |  |  | 35.58 |
| LNS8801 | US073-0000619-O | GPCR | cAMP | Agonist | HTR1A | EC50 | > | 10 |  |  |  | 0 |
| LNS8801 | US073-0000619-O | GPCR | cAMP | Antagonist | HTR1A | IC50 | > | 10 |  |  |  | 32.28 |
| LNS8801 | US073-0000619-O | GPCR | cAMP | Agonist | HTR1B | EC50 | > | 10 |  |  |  | 19.9 |
| LNS8801 | US073-0000619-O | GPCR | cAMP | Antagonist | HTR1B | IC50 | > | 10 |  |  |  | 27.34 |
| LNS8801 | US073-0000619-O | GPCR | Calcium Flux | Agonist | HTR2A | EC50 | > | 10 |  |  |  | 0.86 |
| LNS8801 | US073-0000619-O | GPCR | Calcium Flux | Antagonist | HTR2A | IC50 | > | 10 |  |  |  | 12.79 |
| LNS8801 | US073-0000619-O | GPCR | Calcium Flux | Agonist | HTR2B | EC50 | > | 10 |  |  |  | 1.04 |
| LNS8801 | US073-0000619-O | GPCR | Calcium Flux | Antagonist | HTR2B | IC50 | = | 8.18032 | 1.17 | 0 | 100 | 53.37 |
| LNS8801 | US073-0000619-O | GPCR | cAMP | Agonist | OPRD1 | EC50 | = | 0.87018 | 1.2 | -7.34 | 52.97 | 47.87 |
| LNS8801 | US073-0000619-O | GPCR | cAMP | Antagonist | OPRD1 | IC50 | > | 10 |  |  |  | 7.76 |
| LNS8801 | US073-0000619-O | GPCR | cAMP | Agonist | OPRK1 | EC50 | > | 10 |  |  |  | 12.8 |
| LNS8801 | US073-0000619-O | GPCR | cAMP | Antagonist | OPRK1 | IC50 | > | 10 |  |  |  | 3.8 |
| LNS8801 | US073-0000619-O | GPCR | cAMP | Agonist | OPRM1 | EC50 | > | 10 |  |  |  | 6.68 |
| LNS8801 | US073-0000619-O | GPCR | cAMP | Antagonist | OPRM1 | IC50 | = | 3.54805 | 4.65 | 4.88 | 59.98 | 59.54 |
| LNS8801 | US073-0000619-O | Ion Channel | Ion Channel | Blocker | CAV1.2 | IC50 | > | 10 |  |  |  | 2.65 |
| LNS8801 | US073-0000619-O | Ion Channel | Ion Channel | Blocker | GABAA | IC50 | > | 10 |  |  |  | 15.6 |
| LNS8801 | US073-0000619-O | Ion Channel | Ion Channel | Opener | GABAA | EC50 | > | 10 |  |  |  | 2.5 |
| LNS8801 | US073-0000619-O | Ion Channel | Ion Channel | Blocker | hERG | IC50 | > | 10 |  |  |  | 8.45 |
| LNS8801 | US073-0000619-O | Ion Channel | Ion Channel | Blocker | HTR3A | IC50 | > | 10 |  |  |  | 29.12 |
| LNS8801 | US073-0000619-O | Ion Channel | Ion Channel | Opener | HTR3A | EC50 | > | 10 |  |  |  | 0 |
| LNS8801 | US073-0000619-O | Ion Channel | Ion Channel | Blocker | KvLQT1/minK | IC50 | > | 10 |  |  |  | 28.94 |
| LNS8801 | US073-0000619-O | Ion Channel | Ion Channel | Opener | KvLQT1/minK | EC50 | > | 10 |  |  |  | 1.77 |
| LNS8801 | US073-0000619-O | Ion Channel | Ion Channel | Blocker | nAChR(a4/b2) | IC50 | > | 10 |  |  |  | 20.27 |
| LNS8801 | US073-0000619-O | Ion Channel | Ion Channel | Opener | nAChR(a4/b2) | EC50 | > | 10 |  |  |  | 0 |
| LNS8801 | US073-0000619-O | Ion Channel | Ion Channel | Blocker | NAV1.5 | IC50 | > | 10 |  |  |  | 41.42 |
| LNS8801 | US073-0000619-O | Ion Channel | Ion Channel | Blocker | NMDAR (1A/2B) | IC50 | > | 10 |  |  |  | 0 |
| LNS8801 | US073-0000619-O | Ion Channel | Ion Channel | Opener | NMDAR (1A/2B) | EC50 | > | 10 |  |  |  | 0 |
| LNS8801 | US073-0000619-O | Kinases | Binding | Inhibitor | INSR | IC50 | > | 10 |  |  |  | 5.77 |
| LNS8801 | US073-0000619-O | Kinases | Binding | Inhibitor | LCK | IC50 | > | 10 |  |  |  | 0 |
| LNS8801 | US073-0000619-O | Kinases | Binding | Inhibitor | ROCK1 | IC50 | > | 10 |  |  |  | 31.82 |
| LNS8801 | US073-0000619-O | Kinases | Binding | Inhibitor | VEGFR2 | IC50 | > | 10 |  |  |  | 1.13 |
| LNS8801 | US073-0000619-O | NHR | NHR Nuclear Translocation | Agonist | AR | EC50 | > | 10 |  |  |  | 0.06 |
| LNS8801 | US073-0000619-O | NHR | NHR Nuclear Translocation | Antagonist | AR | IC50 | > | 10 |  |  |  | 0 |
| LNS8801 | US073-0000619-O | NHR | NHR Protein Interaction | Agonist | GR | EC50 | > | 10 |  |  |  | 0.21 |
| LNS8801 | US073-0000619-O | NHR | NHR Protein Interaction | Antagonist | GR | IC50 | > | 10 |  |  |  | 22.45 |
| LNS8801 | US073-0000619-O | Non-Kinase Enzymes | Enzymatic | Inhibitor | AChE | IC50 | > | 10 |  |  |  | 12.38 |
| LNS8801 | US073-0000619-O | Non-Kinase Enzymes | Enzymatic | Inhibitor | COX1 | IC50 | > | 10 |  |  |  | 9.95 |
| LNS8801 | US073-0000619-O | Non-Kinase Enzymes | Enzymatic | Inhibitor | COX2 | IC50 | > | 10 |  |  |  | 10.96 |
| LNS8801 | US073-0000619-O | Non-Kinase Enzymes | Enzymatic | Inhibitor | MAOA | IC50 | > | 10 |  |  |  | 0 |
| LNS8801 | US073-0000619-O | Non-Kinase Enzymes | Enzymatic | Inhibitor | PDE3A | IC50 | > | 10 |  |  |  | 5.8 |
| LNS8801 | US073-0000619-O | Non-Kinase Enzymes | Enzymatic | Inhibitor | PDE4D2 | IC50 | > | 10 |  |  |  | 11.19 |
| LNS8801 | US073-0000619-O | Transporter | Transporter | Blocker | DAT | IC50 | > | 10 |  |  |  | 0 |
| LNS8801 | US073-0000619-O | Transporter | Transporter | Blocker | NET | IC50 | > | 10 |  |  |  | 0 |
| LNS8801 | US073-0000619-O | Transporter | Transporter | Blocker | SERT | IC50 | > | 10 |  |  |  | 0 |
| LNS8812 | US073-0000619-O | GPCR | Calcium Flux | Agonist | ADORA2A | EC50 | > | 10 |  |  |  | 0 |
| LNS8812 | US073-0000619-O | GPCR | Calcium Flux | Antagonist | ADORA2A | IC50 | > | 10 |  |  |  | 0 |
| LNS8812 | US073-0000619-O | GPCR | Calcium Flux | Agonist | ADRA1A | EC50 | > | 10 |  |  |  | 0 |
| LNS8812 | US073-0000619-O | GPCR | Calcium Flux | Antagonist | ADRA1A | IC50 | > | 10 |  |  |  | 0 |
| LNS8812 | US073-0000619-O | GPCR | cAMP | Agonist | ADRA2A | EC50 | > | 10 |  |  |  | 0 |
| LNS8812 | US073-0000619-O | GPCR | cAMP | Antagonist | ADRA2A | IC50 | = | 2.07253 | 2.45 | 0 | 59.26 | 58.17 |
| LNS8812 | US073-0000619-O | GPCR | cAMP | Agonist | ADRB1 | EC50 | > | 10 |  |  |  | 1.52 |
| LNS8812 | US073-0000619-O | GPCR | cAMP | Antagonist | ADRB1 | IC50 | > | 10 |  |  |  | 28.15 |
| LNS8812 | US073-0000619-O | GPCR | cAMP | Agonist | ADRB2 | EC50 | > | 10 |  |  |  | 1.68 |
| LNS8812 | US073-0000619-O | GPCR | cAMP | Antagonist | ADRB2 | IC50 | > | 10 |  |  |  | 3.47 |
| LNS8812 | US073-0000619-O | GPCR | Calcium Flux | Agonist | AVPR1A | EC50 | > | 10 |  |  |  | 0 |
| LNS8812 | US073-0000619-O | GPCR | Calcium Flux | Antagonist | AVPR1A | IC50 | > | 10 |  |  |  | 0 |
| LNS8812 | US073-0000619-O | GPCR | Calcium Flux | Agonist | CCKAR | EC50 | > | 10 |  |  |  | 1.15 |
| LNS8812 | US073-0000619-O | GPCR | Calcium Flux | Antagonist | CCKAR | IC50 | > | 10 |  |  |  | 0 |
| LNS8812 | US073-0000619-O | GPCR | Calcium Flux | Agonist | CHRM1 | EC50 | > | 10 |  |  |  | 0 |
| LNS8812 | US073-0000619-O | GPCR | Calcium Flux | Antagonist | CHRM1 | IC50 | > | 10 |  |  |  | 0 |
| LNS8812 | US073-0000619-O | GPCR | cAMP | Agonist | CHRM2 | EC50 | > | 10 |  |  |  | 0 |
| LNS8812 | US073-0000619-O | GPCR | cAMP | Antagonist | CHRM2 | IC50 | > | 10 |  |  |  | 32.13 |
| LNS8812 | US073-0000619-O | GPCR | Calcium Flux | Agonist | CHRM3 | EC50 | > | 10 |  |  |  | 2.64 |
| LNS8812 | US073-0000619-O | GPCR | Calcium Flux | Antagonist | CHRM3 | IC50 | > | 10 |  |  |  | 0 |
| LNS8812 | US073-0000619-O | GPCR | cAMP | Agonist | CNR1 | EC50 | > | 10 |  |  |  | 0 |
| LNS8812 | US073-0000619-O | GPCR | cAMP | Antagonist | CNR1 | IC50 | = | 3.13957 | 1.5 | 0 | 100 | 84.54 |
| LNS8812 | US073-0000619-O | GPCR | cAMP | Agonist | CNR2 | EC50 | > | 10 |  |  |  | 0 |
| LNS8812 | US073-0000619-O | GPCR | cAMP | Antagonist | CNR2 | IC50 | > | 10 |  |  |  | 25.96 |
| LNS8812 | US073-0000619-O | GPCR | cAMP | Agonist | DRD1 | EC50 | > | 10 |  |  |  | 4.79 |
| LNS8812 | US073-0000619-O | GPCR | cAMP | Antagonist | DRD1 | IC50 | > | 10 |  |  |  | 18.52 |
| LNS8812 | US073-0000619-O | GPCR | cAMP | Agonist | DRD2S | EC50 | > | 10 |  |  |  | 0 |
| LNS8812 | US073-0000619-O | GPCR | cAMP | Antagonist | DRD2S | IC50 | > | 10 |  |  |  | 0 |
| LNS8812 | US073-0000619-O | GPCR | Calcium Flux | Agonist | EDNRA | EC50 | > | 10 |  |  |  | 0 |
| LNS8812 | US073-0000619-O | GPCR | Calcium Flux | Antagonist | EDNRA | IC50 | > | 10 |  |  |  | 0 |
| LNS8812 | US073-0000619-O | GPCR | Calcium Flux | Agonist | HRH1 | EC50 | > | 10 |  |  |  | 1.95 |
| LNS8812 | US073-0000619-O | GPCR | Calcium Flux | Antagonist | HRH1 | IC50 | > | 10 |  |  |  | 4.63 |
| LNS8812 | US073-0000619-O | GPCR | cAMP | Agonist | HRH2 | EC50 | > | 10 |  |  |  | 3.46 |
| LNS8812 | US073-0000619-O | GPCR | cAMP | Antagonist | HRH2 | IC50 | > | 10 |  |  |  | 14.84 |
| LNS8812 | US073-0000619-O | GPCR | cAMP | Agonist | HTR1A | EC50 | > | 10 |  |  |  | 0 |
| LNS8812 | US073-0000619-O | GPCR | cAMP | Antagonist | HTR1A | IC50 | = | 2.1564 | 1.5 | 12.53 | 46.39 | 43.33 |
| LNS8812 | US073-0000619-O | GPCR | cAMP | Agonist | HTR1B | EC50 | > | 10 |  |  |  | 3.51 |
| LNS8812 | US073-0000619-O | GPCR | cAMP | Antagonist | HTR1B | IC50 | > | 10 |  |  |  | 33.33 |
| LNS8812 | US073-0000619-O | GPCR | Calcium Flux | Agonist | HTR2A | EC50 | > | 10 |  |  |  | 0 |
| LNS8812 | US073-0000619-O | GPCR | Calcium Flux | Antagonist | HTR2A | IC50 | > | 10 |  |  |  | 1.5 |
| LNS8812 | US073-0000619-O | GPCR | Calcium Flux | Agonist | HTR2B | EC50 | > | 10 |  |  |  | 0.49 |
| LNS8812 | US073-0000619-O | GPCR | Calcium Flux | Antagonist | HTR2B | IC50 | > | 10 |  |  |  | 0 |
| LNS8812 | US073-0000619-O | GPCR | cAMP | Agonist | OPRD1 | EC50 | > | 10 |  |  |  | 30.98 |
| LNS8812 | US073-0000619-O | GPCR | cAMP | Antagonist | OPRD1 | IC50 | > | 10 |  |  |  | 0 |
| LNS8812 | US073-0000619-O | GPCR | cAMP | Agonist | OPRK1 | EC50 | > | 10 |  |  |  | 0 |
| LNS8812 | US073-0000619-O | GPCR | cAMP | Antagonist | OPRK1 | IC50 | > | 10 |  |  |  | 5.02 |
| LNS8812 | US073-0000619-O | GPCR | cAMP | Agonist | OPRM1 | EC50 | > | 10 |  |  |  | 0 |
| LNS8812 | US073-0000619-O | GPCR | cAMP | Antagonist | OPRM1 | IC50 | > | 10 |  |  |  | 2.61 |
| LNS8812 | US073-0000619-O | Ion Channel | Ion Channel | Blocker | CAV1.2 | IC50 | > | 10 |  |  |  | 19.41 |
| LNS8812 | US073-0000619-O | Ion Channel | Ion Channel | Blocker | GABAA | IC50 | > | 10 |  |  |  | 10.51 |
| LNS8812 | US073-0000619-O | Ion Channel | Ion Channel | Opener | GABAA | EC50 | > | 10 |  |  |  | 10.11 |
| LNS8812 | US073-0000619-O | Ion Channel | Ion Channel | Blocker | hERG | IC50 | > | 10 |  |  |  | 11.68 |

| Compound Name | Order ID | Target Class | Assay Name | Mode | Assay Target | Result Type | Value Prefix | RC50 (µM) | Hill | Curve Bottom | Curve Top | Max Response |
| --- | --- | --- | --- | --- | --- | --- | --- | --- | --- | --- | --- | --- |
| LNS8812 | US073-0000619-O | Ion Channel | Ion Channel | Blocker | HTR3A | IC50 | > | 10 |  |  |  | 40.16 |
| LNS8812 | US073-0000619-O | Ion Channel | Ion Channel | Opener | HTR3A | EC50 | > | 10 |  |  |  | 0.46 |
| LNS8812 | US073-0000619-O | Ion Channel | Ion Channel | Blocker | KvLQT1/minK | IC50 | > | 10 |  |  |  | 56.4 |
| LNS8812 | US073-0000619-O | Ion Channel | Ion Channel | Opener | KvLQT1/minK | EC50 | > | 10 |  |  |  | 0 |
| LNS8812 | US073-0000619-O | Ion Channel | Ion Channel | Blocker | nAChR(a4/b2) | IC50 | > | 10 |  |  |  | 42.99 |
| LNS8812 | US073-0000619-O | Ion Channel | Ion Channel | Opener | nAChR(a4/b2) | EC50 | > | 10 |  |  |  | 12.7 |
| LNS8812 | US073-0000619-O | Ion Channel | Ion Channel | Blocker | NAV1.5 | IC50 | > | 10 |  |  |  | 29.44 |
| LNS8812 | US073-0000619-O | Ion Channel | Ion Channel | Blocker | NMDAR (1A/2B) | IC50 | > | 10 |  |  |  | 0 |
| LNS8812 | US073-0000619-O | Ion Channel | Ion Channel | Opener | NMDAR (1A/2B) | EC50 | > | 10 |  |  |  | 0 |
| LNS8812 | US073-0000619-O | Kinases | Binding | Inhibitor | INSR | IC50 | > | 10 |  |  |  | 22.51 |
| LNS8812 | US073-0000619-O | Kinases | Binding | Inhibitor | LCK | IC50 | > | 10 |  |  |  | 14 |
| LNS8812 | US073-0000619-O | Kinases | Binding | Inhibitor | ROCK1 | IC50 | > | 10 |  |  |  | 30.33 |
| LNS8812 | US073-0000619-O | Kinases | Binding | Inhibitor | VEGFR2 | IC50 | > | 10 |  |  |  | 0 |
| LNS8812 | US073-0000619-O | NHR | NHR Nuclear Translocation | Agonist | AR | EC50 | > | 10 |  |  |  | 0 |
| LNS8812 | US073-0000619-O | NHR | NHR Nuclear Translocation | Antagonist | AR | IC50 | = | 4.76277 | 1.22 | 0 | 100 | 65.49 |
| LNS8812 | US073-0000619-O | NHR | NHR Protein Interaction | Agonist | GR | EC50 | > | 10 |  |  |  | 0.21 |
| LNS8812 | US073-0000619-O | NHR | NHR Protein Interaction | Antagonist | GR | IC50 | > | 10 |  |  |  | 16.74 |
| LNS8812 | US073-0000619-O | Non-Kinase Enzymes | Enzymatic | Inhibitor | AChE | IC50 | > | 10 |  |  |  | 8.34 |
| LNS8812 | US073-0000619-O | Non-Kinase Enzymes | Enzymatic | Inhibitor | COX1 | IC50 | > | 10 |  |  |  | 2.06 |
| LNS8812 | US073-0000619-O | Non-Kinase Enzymes | Enzymatic | Inhibitor | COX2 | IC50 | > | 10 |  |  |  | 0 |
| LNS8812 | US073-0000619-O | Non-Kinase Enzymes | Enzymatic | Inhibitor | MAOA | IC50 | > | 10 |  |  |  | 0 |
| LNS8812 | US073-0000619-O | Non-Kinase Enzymes | Enzymatic | Inhibitor | PDE3A | IC50 | > | 10 |  |  |  | 0.91 |
| LNS8812 | US073-0000619-O | Non-Kinase Enzymes | Enzymatic | Inhibitor | PDE4D2 | IC50 | > | 10 |  |  |  | 6.54 |
| LNS8812 | US073-0000619-O | Transporter | Transporter | Blocker | DAT | IC50 | > | 10 |  |  |  | 0 |
| LNS8812 | US073-0000619-O | Transporter | Transporter | Blocker | NET | IC50 | > | 10 |  |  |  | 0 |
| LNS8812 | US073-0000619-O | Transporter | Transporter | Blocker | SERT | IC50 | > | 10 |  |  |  | 0 |

